# Mapping early PRC2 nucleation sites upon Suz12 reintroduction reveals features of *de novo* Polycomb recruitment

**DOI:** 10.1101/2025.09.02.673664

**Authors:** Itzel Alejandra Hernández-Romero, Carlos Alberto Peralta-Alvarez, Abraham Román-Figueroa, Nallely Cano-Domínguez, Maribel Soto-Nava, Hongwei Zhou, Xin Huang, Félix Recillas-Targa, Augusto Cesar Poot-Hernandez, Mayra Furlan-Magaril, Santiago Avila-Rios, David Valle-Garcia, Jianlong Wang, Victor Julian Valdes

## Abstract

Polycomb domains safeguard cell identity by maintaining lineage-specific chromatin states enriched in repressive histone modifications, preserving the epigenetic memory of cell lineages. While Polycomb Repressive Complex 2 (PRC2) can re-establish its occupancy after perturbation, the mechanisms that guide *de novo* Polycomb recruitment remain unclear. To address this, we engineered an auxin-inducible degradation system to reversibly deplete and reintroduce the endogenous PRC2 core subunit Suz12 in mouse embryonic stem cells (mESCs). Genome-wide profiling at an early recovery time point revealed ∼1,100 PRC2 nucleation sites, characterized by rapid Suz12 and histone H3K27me3 re-accumulation with strong signal, with minimal impact on gene expression. These sites were significantly enriched at bivalent promoters, coinciding with unmethylated CpG islands and chromatin states associated with developmental regulation, and were largely conserved in differentiated cells. Motif analysis identified G/C-rich DNA sequences associated with E2F and zinc-finger proteins, alongside strong co-occupancy with MTF2 and JARID2, two PRC2 cofactors previously implicated in Polycomb targeting. Notably, a subset of nucleation sites overlapped with long-range chromatin interaction anchors in histone H3K27me3 HiChIP datasets. These findings reveal that PRC2 *de novo* nucleation sites are associated with a combination of chromatin states, DNA sequence features, cofactor co-occupancy and spatial genome organization, suggesting that epigenetic memory can be re-established through defined genomic and chromatin features.

**Author summary:** Polycomb group proteins are key epigenetic regulators that silence gene expression by establishing and dispersing repressive chromatin domains marked by histone modifications such as histone H3K27me3 and H2AK119ub1 and are critical for defining cell identity. During differentiation, Polycomb domains are dynamically redistributed, implying a mechanism for *de novo* targeting to specific loci. How the Polycomb Repressive Complex 2 (PRC2) is initially recruited to these nucleation sites and which features stabilize its binding remain poorly understood. To explore this, we studied the characteristics of the *de novo* recruitment sites of PRC2 in mouse embryonic stem cells (mESCs) using an auxin-inducible degradation (AID) system targeting PRC2 to deplete and reintroduce the core subunit Suz12. We identified recruitment sites after complete clearance of the histone H3K27me3 through ChIP-seq at a very early time point after Suz12 reintroduction. Most nucleation sites were located at bivalent promoters of developmental genes and correlated with unmethylated CpG islands. Motif analysis revealed over-represented sequences and accessory partners such as MTF2 and JARID2, along with long-range chromatin interactions. These nucleation sites were also conserved in differentiated cells, highlighting their potential role in developmental regulation. Our study provides insight into how Polycomb domains are established and how epigenetic memory is maintained in stem cells.

## Introduction

The Polycomb Repressive Complex 2 (PRC2) is a key epigenetic regulator that fine-tunes gene expression through the deposition of tri-methylation of histone H3 at lysine 27 (H3K27me3), which is associated with gene silencing (1,2). This histone posttranslational modification helps to establish facultative heterochromatin regions, characterized by low transcriptional rate (3,4). Like gene expression patterns, H3K27me3 chromatin domains are highly cell-type specific (5). Classical examples include PRC2-mediated transcriptional repression of somatic genes in embryonic stem cells (ESCs), while conversely repressing the expression of pluripotency genes in differentiated cells (6). Therefore, understanding the dynamic recruitment of Polycomb to chromatin remains a central question, as impairment of PRC2 is associated with defects in embryonic development and the appearance of diseases such as cancer (7,8).

During DNA replication, parental histones carrying histone H3K27me3 are recycled to enable the Polycomb machinery to restore local repressive regions (9). After that, two main positive feedback axes reinforce Polycomb recruitment: (i) recognition and catalysis of histone H3K27me3 by PRC2 itself (10), and (ii) the binding of PRC2 to the Polycomb Repressive Complex 1 (PRC1) histone mark (H2AK119ub1) (11). However, the specific mechanisms by which Polycomb identifies its target loci and generates new repressive domains — particularly following domain erosion or during differentiation — remain poorly understood (12,13).

Recent findings have proposed *de novo* recruitment as an additional mechanism that operates independently of the positive feedback of pre-existing histone marks (14–17). Although unmethylated CpG islands (CGI) are often enriched at these sites (18), CGI content alone cannot explain the diversity and cell-type specificity of Polycomb domains. Additional factors such as *cis*-regulatory elements, transcription factor interactions, or epigenetic crosstalk may act as nucleation centers from which H3K27me3 domains disperse, feeding back into the steady-state.

Prolonged depletion and subsequent reintroduction of Polycomb group (PcG) proteins have enabled the dissection of Polycomb’s ability to accurately reconstitute lineage-specific chromatin patterns, even in the absence of pre-existing repressive histone marks (14–17). For example, ablation of the PRC2 subunit Suz12 erodes the mark, but its re-expression restores the histone H3K27me3 pattern at CGI (15). Similarly, reintroduction of EED unable to spread the repressive domains leads to spatial stalling of PRC2 at nucleation sites, forming clusters (14). Additionally, reintroduction of the catalytic subunit EZH2 in Ezh1/Ezh2 double knockouts leads to restoration of histone H3K27me3 levels and PRC2 occupancy (16). While these studies focused on steady-state profiles, they collectively suggest that PRC2 recruitment mechanisms can operate intrinsically, even in the absence of the histone H3K27me3 signal. However, it remains unclear how *de novo* sites are identified in a developmental and physiological context, and what actors stabilize PRC2 at the domain nucleation sites.

The ability of PRC2 to locate its targets depends on its core subunit Suz12. Suz12 knockout assays have demonstrated that the patterns of histone H3K27 methylation can only be restored upon reintroducing Suz12 (15). This subunit serves as a structural platform that coordinates both the assembly of catalytic core subunits and accessory proteins essential for chromatin recruitment, as none of the core PRC2 subunits possess intrinsic DNA-binding ability (9,19–23). Suz12 possesses two functional components: the VEFS domain that mediates the interaction with other PRC2 members, and the N-terminal region, which couples accessory subunits involved in chromatin binding (1,22). Notably, Suz12 recruitment can occur without pre-existing H3K27me3 or H2AK119ub1, indicating histone mark independent targeting (17).

To investigate the early events of *de novo* PRC2 recruitment, we implemented an auxin-inducible and reversible system targeting the endogenous Suz12 in mouse embryonic stem cells (mESCs). Auxin treatment led to the rapid degradation of Suz12 and consequent erosion of H3K27me3 domains. Following reintroduction of Suz12, we performed Chromatin Immunoprecipitation (ChIP-seq) at a very early time point to map nucleation sites before domain dispersal. Unlike prior studies focused on restored steady-state profiles, our approach captures the earliest stages, revealing that nucleation sites are significantly enriched at bivalent promoters, display higher histone H3K27me3 re-accumulation, overlap with MTF2 and JARID2 cofactors, and coincide with unmethylated CGI and long-range H3K27me3 chromatin loops. This study provides insights into the initial steps that govern PRC2 recruitment and histone H3K27me3 distribution, providing a multidimensional view of early Polycomb recruitment in the context of epigenetic memory.

## Results

### Establishment of an Auxin-inducible Suz12 depletion system to capture *de novo* PRC2 recruitment events

Mouse embryonic stem cells (mESCs) retain features of the preimplantation epiblast, including naïve pluripotency and self-renewal capability, and importantly, they can proliferate in the absence of PRC2 (24,25). To investigate early PRC2 recruitment, we generated a homozygous mESCs line incorporating an Auxin Inducible and Degradation system (AID) targeting the endogenous Suz12 using CRISPR-Cas9, along with the introduction of the OsTIR1 receptor (26) (Fig 1A and S1A-G). In this system, auxin induces rapid degradation of the transgenic protein (also targeted with a fluorescent protein), while auxin removal leads to Suz12 re-accumulation (Fig 1B).

**Fig 1.**
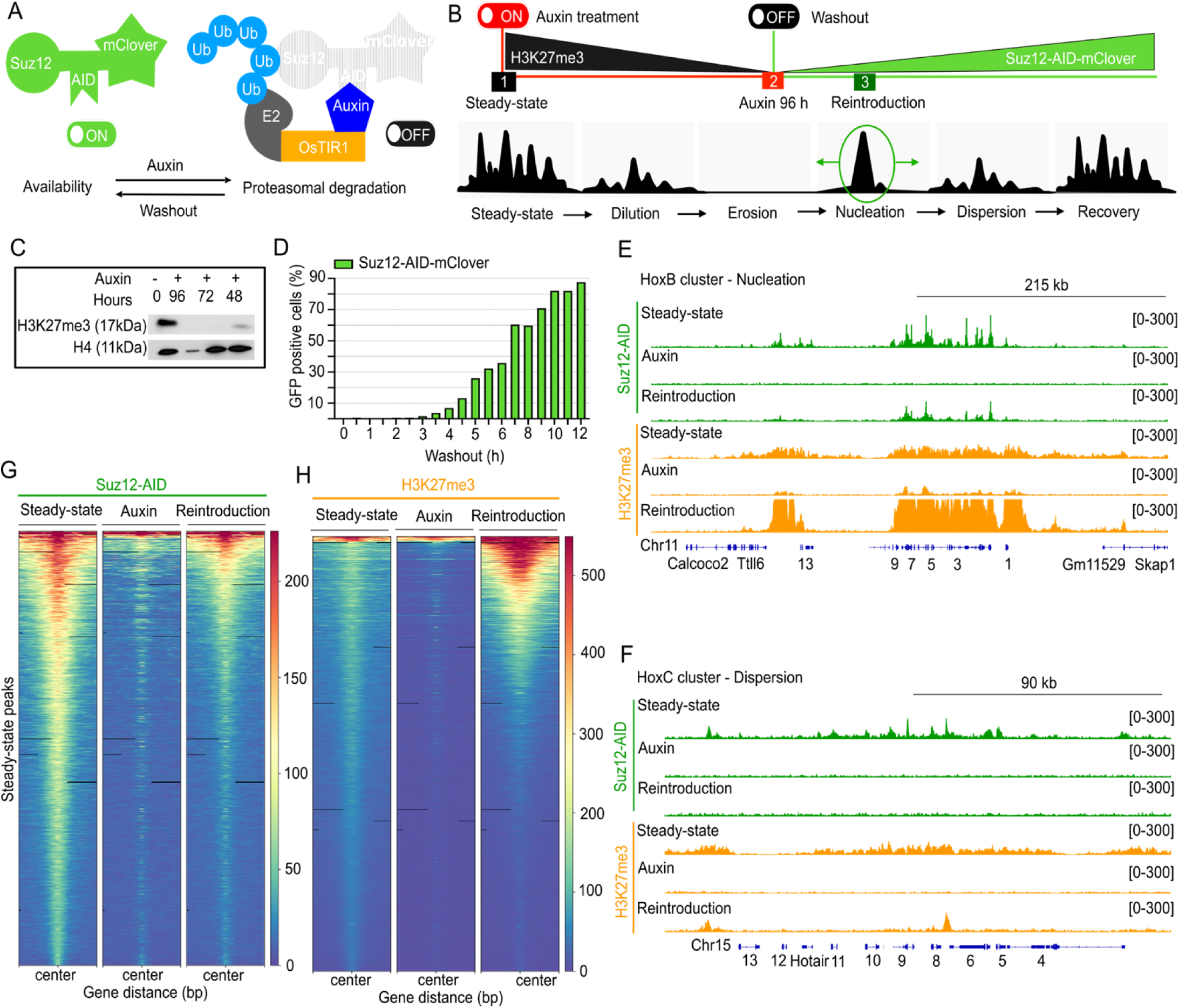
ON/OFF system for Suz12. (**A)** Auxin Inducible-Degradation (AID) system of Suz12 in mouse embryonic stem cells (mESCs). The endogenous Suz12 Open Reading Frame (ORF) was tagged with an AID-mClover cassette via CRISPR-Cas9 in cells expressing an OsTIR1 transgene, enabling auxin (IAA)-mediated proteasomal degradation. **(B)** Experimental design for PRC2 domain erosion and Suz12 *de novo* recruitment assay: (1) *Steady-state*; (2) *Auxin* treatment for Suz12 degradation and histone H3K27me3 loss; (3) *Reintroduction:* auxin removal and re-accumulation of Suz12. **(C)** Western blot showing progressive histone H3K27me3 loss at different auxin treatment time points. **(D)** Time course of Suz12-mClover recovery after auxin washout, monitored by FACS over 12 hours. At 4.5 hours, Suz12 reaches 11.5% of GFP+ cells. **(E-F)** ChIP-seq tracks for Suz12 (green) and histone H3K27me3 (orange) 4.5 hours after auxin removal at the representative loci, including nucleation sites at the HoxB cluster (E) and the HoxC cluster that shows signal dispersion (F). **(G-H)** Global ChIP-seq signal for Suz12 (G) and histone H3K27me3 (H) at *Control-exclusive* peaks through the three conditions. Heatmaps of read density are within a ±1.5 kb centered on the maximum value of the peak signal.

To identify *de novo* recruitment sites independent of histone H3K27me3, we treated cells with auxin for 96 hours (∼6 cell divisions (27)) to allow sustained absence of PRC2 and depletion of histone H3K27me3 via dilution during replication (Fig 1C) (28). At this time, 96.4% of the cells were arrested in G1 phase (Fig S1H), likely due to derepression of cell cycle regulators (e.g., cyclins D1/E1, Ink4a locus) (29). Cells resumed S/G2 phase progression upon auxin washout. To capture early recruitment events before domain spreading, we calibrated the recovery time to 4.5 hours post-washout, corresponding to ∼10% of the Suz12 fluorescent signal (Fig 1D, S1I), at which we performed ChIP-seq for histone H3K27me3 and Suz12. Inspection of the ChIP-seq enrichment signals at two Hox gene clusters validated our system. While the treatment depleted Suz12 and histone H3K27me3, upon *Reintroduction* with auxin removal, the signals were recovered at the nucleation site within the HoxB cluster but not in the distal HoxC region (Fig 1E and 1F), consistent with prior report (14).

Genome-wide ChIP-seq analysis confirmed global loss of Suz12 and histone H3K27me3 upon auxin treatment, followed by partial restoration during *Reintroduction* (4.5 hours after auxin washout) (Fig 1G and 1H). We interpreted this as early *de nov*o recruitment of Suz12 and re-establishment of histone H3K27me3 at pre-perturbation sites. Interestingly, we noticed an increased deposition of histone H3K27me3 at nucleation sites during *Reintroduction*, which was not observed at dispersal sites. This effect may reflect preferential deposition of histone H3K27me3 at nucleation sites during early *de novo* recruitment. Consistent with this, prior work reported higher local catalytic rates of H3K27 methylation at putative nucleation sites (1) and classified them as “strong” or “weak” based on H3K27me3 deposition (14). Overall, our AID-Suz12 ON/OFF system enables time-resolved capture of early PRC2 recruitment events before domain spreading.

### Genome-wide identification and classification of PRC2 *de novo* nucleation sites

A detailed analysis of all peaks across the three conditions (*Steady-state, Auxin, and Reintroduction)* revealed that auxin treatment led to a significant decrease in Suz12 binding, with a reduction of almost 85% of its target sites lost (Fig 2A). We detected a modest relocalization of Suz12 to new targets in *Auxin* and *Reintroduction* conditions (372 and 448 peaks, respectively), which were absent in *Steady-state* (*Control*). These likely reflect transient or opportunistic binding under perturbed chromatin states (30,31), even in controlled culture conditions that preserve pluripotency as the 2i media. The new targets are mainly intergenic and intronic with no distinctive features detected. Additionally, a small subset of 896 Suz12 peaks persisted under *Auxin* treatment. These peaks were largely intergenic and lacked consistent chromatin or transcriptional features. In order to study the erasure and restoration of Suz12 from the *Steady-state*, we excluded newly acquired peaks from further analysis.

**Fig 2.**
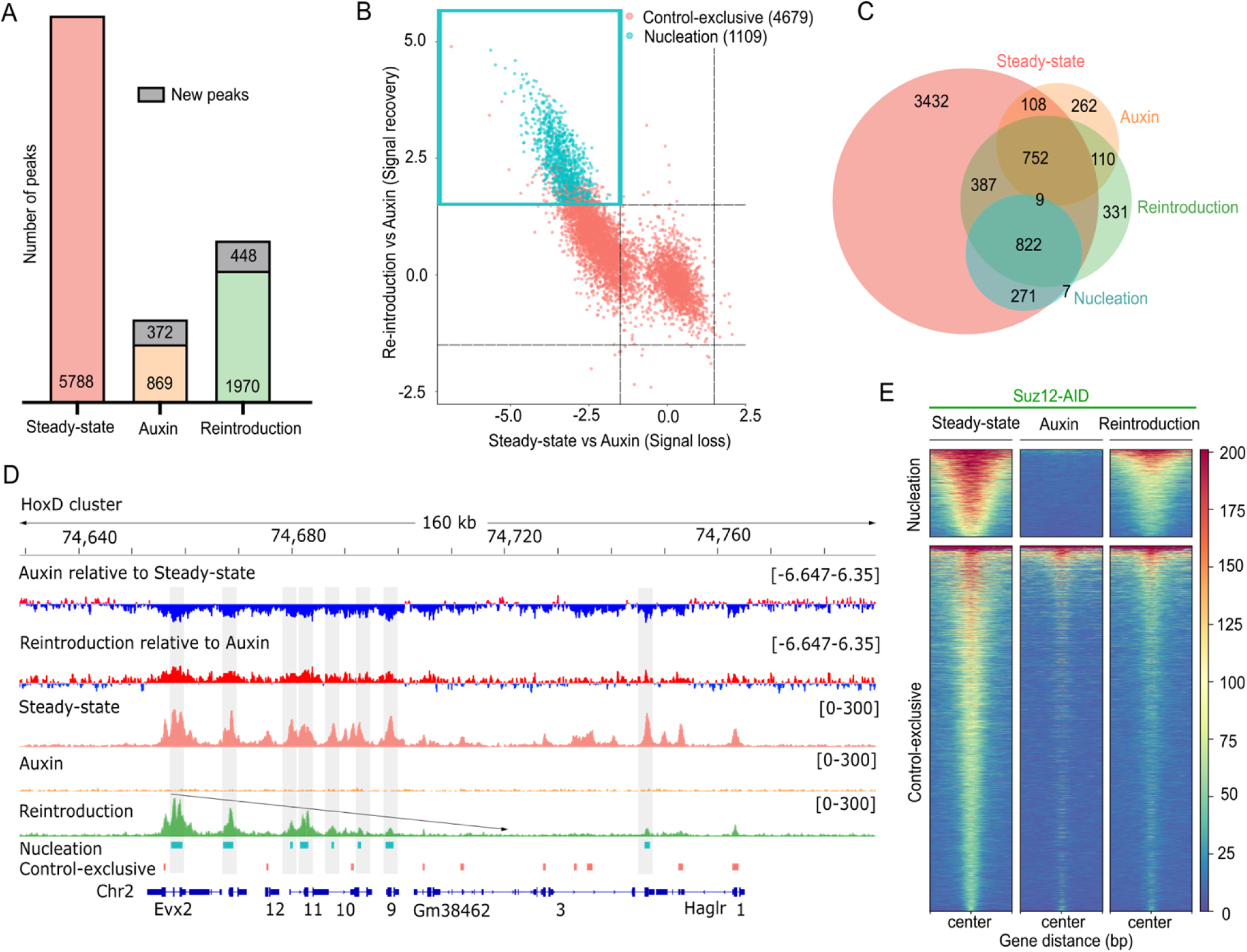
Auxin-inducible degradation of Suz12 enables identification of *de novo* nucleation sites. **(A)** Total number of Suz12 peaks detected across the three conditions. Bar plots indicate peak counts per category, including persistent peaks during *Auxin*, and newly acquired peaks (gray). **(B)** Identification of Suz12 nucleation sites based on dynamic signal changes upon recovery. Each point represents a Suz12 ChIP-seq peak plotted by its fold change upon recovery or loss, versus *Auxin*. The quadrant highlights the peaks that experienced a signal loss with *Auxin* but recovered upon washout (*Reintroduction*) and presumably nucleation true sites. Statistically significant peaks are shown in blue (|log2FoldChange| > 1.5, adjusted P value < 0.005). **(C)** Peak overlap summary across datasets. **(D)** ChIP-seq tracks for Suz12 after auxin removal at representative loci: the Evx2 nucleation site within the HoxD cluster. Tracks signals for *Auxin* relative to *Steady-state* and *Reintroduction* relative to *Auxin* (upper; positive signal in red, negative signal in blue). Nucleation and *Control-exclusive* coordinates (bottom). The diagonal arrow indicates apparent spreading direction from the Evx2 nucleation site across the HoxD cluster. The gray boxes point out the nucleation peaks. **(E)** Heatmaps showing normalized Suz12 ChIP-seq signal intensity across the three conditions at classified nucleation and *Control-exclusive* peaks. Each heatmap is centered on the peak summit within a ±1.5 kb window.

To identify *bona fide* nucleation sites, we normalized Suz12 ChIP-seq read counts to estimate the fold change over each peak using DESeq2, (|log_2_FoldChange| > 1.5, adjusted P-value < 0.005). To characterize the binding dynamic of Suz12 under different conditions, we compared signal intensities at each peak between *Auxin* and *Steady-state*, versus *Reintroduction* and *Auxin*, respectively (Fig 2B). This approach allows us to classify nucleation sites as peaks whose signal dynamics met the following criteria: (i) a significantly decreased signal upon *Auxin* treatment compared to the *Steady-state*, and (ii) a significantly increased signal in the *Reintroduction* compared to the *Auxin* (blue quadrant, Fig 2B). Our strategy identified 1,109 Suz12 nucleation sites (corresponding to 19% of the total *Steady-state* peaks) For analysis purposes, we referred to the rest of the peaks that do not meet the nucleation criteria as “*Control-exclusive*”. Applying the same classification to histone H3K27me3 ChIP-seq data identified 437 nucleation sites (Fig S2A). However, we focused on Suz12, as its binding marks the earliest step in PRC2 recruitment, preceding histone H3K27me3 deposition, and therefore offers a more direct measure of nucleation independent of downstream catalytic activity.

Our intensity-based classification method was both effective and efficient, as it identified most of the putative nucleation sites described in the literature, including sites at genes such as Cyp26b1, Emx1, Evx2, Lhx2, Lmx1b, and Tox2 (14), all of which fell within the first quadrant of our analysis (Fig S2B). We also compared our nucleation set with an independent published dataset that reintroduced an EED-cage mutant (Phe97 or Tyr365 substitution with alanine) unable to spread H3K27me3 domains (14). Remarkably, 908 (81%) of our *de novo* Suz12 peaks overlapped with the 5,128 PRC2 peaks from the EED-cage mutant ChIP-seq dataset (Fig S2C). This suggests our AID-based approach may capture a more stringent set of early, high-confidence PRC2 recruitment events.

To further support this classification, we assessed the overlap of our nucleation peaks with peaks independently called in each condition. As shown in Fig 2C, our nucleation peaks overlapped strongly with those detected upon Suz12 reintroduction, confirming that our approach effectively captures early PRC2 recruitment events that are absent upon auxin treatment but reappear after washout.

We analyze the signal dynamics of nucleation and *Control-exclusive* peaks at the Evx2 nucleation site, which spreads across the HoxD cluster (14) (Fig 2D). The first two tracks illustrate the positive (red) and negative (blue) changes occurring in the *Auxin* compared to the *Steady-state*, as well as the changes after *Reintroduction* compared to the *Auxin* condition. We observed a similar behavior indicating a potential spreading direction for Pax2 and Lbx1 (Fig S2D). Our findings confirm that the nucleation sites exhibit greater intensity changes than the *Control-exclusive* peaks, which are not present at our early *Reintroduction* time point.

We next analyzed whether nucleation sites differ in their size. We found that nucleation sites were consistently broader, with a median size of 1342 bp, whereas *Control-exclusive* sites had a median size of 714 bp (Fig S2E). This is an intriguing finding, especially when we consider the intricate nature of Polycomb domains, which vary from 1 kb to over 100 kb (32). To validate our classification, we separately examined Suz12 (Fig 2E) and histone H3K27me3 (Fig S2F-G) global signals at the nucleation and *Control-exclusive* sites. These visualizations confirmed distinct occupancy patterns that met our selection criteria. Taken together, our analysis demonstrates that the AID-Suz12 system effectively reveals early PRC2 nucleation sites that overcome histone H3K27me3 loss and can rediscover their targets.

### Nucleation sites localize to bivalent promoters and poised chromatin states

We assessed whether our mESCs PRC2-nucleation sites are preserved in differentiated cells. To address this, we re-analyzed publicly available Suz12 ChIP-seq datasets from thymocytes (33), intestinal epithelium (34), and neural progenitor cells (35). Approximately, 70% of nucleation sites overlapped with the 3,512 PRC2 peaks shared across these three lineages (Fig 3A), indicating that most nucleation regions are maintained across tissues derived from the three germ layers.

**Fig 3.**
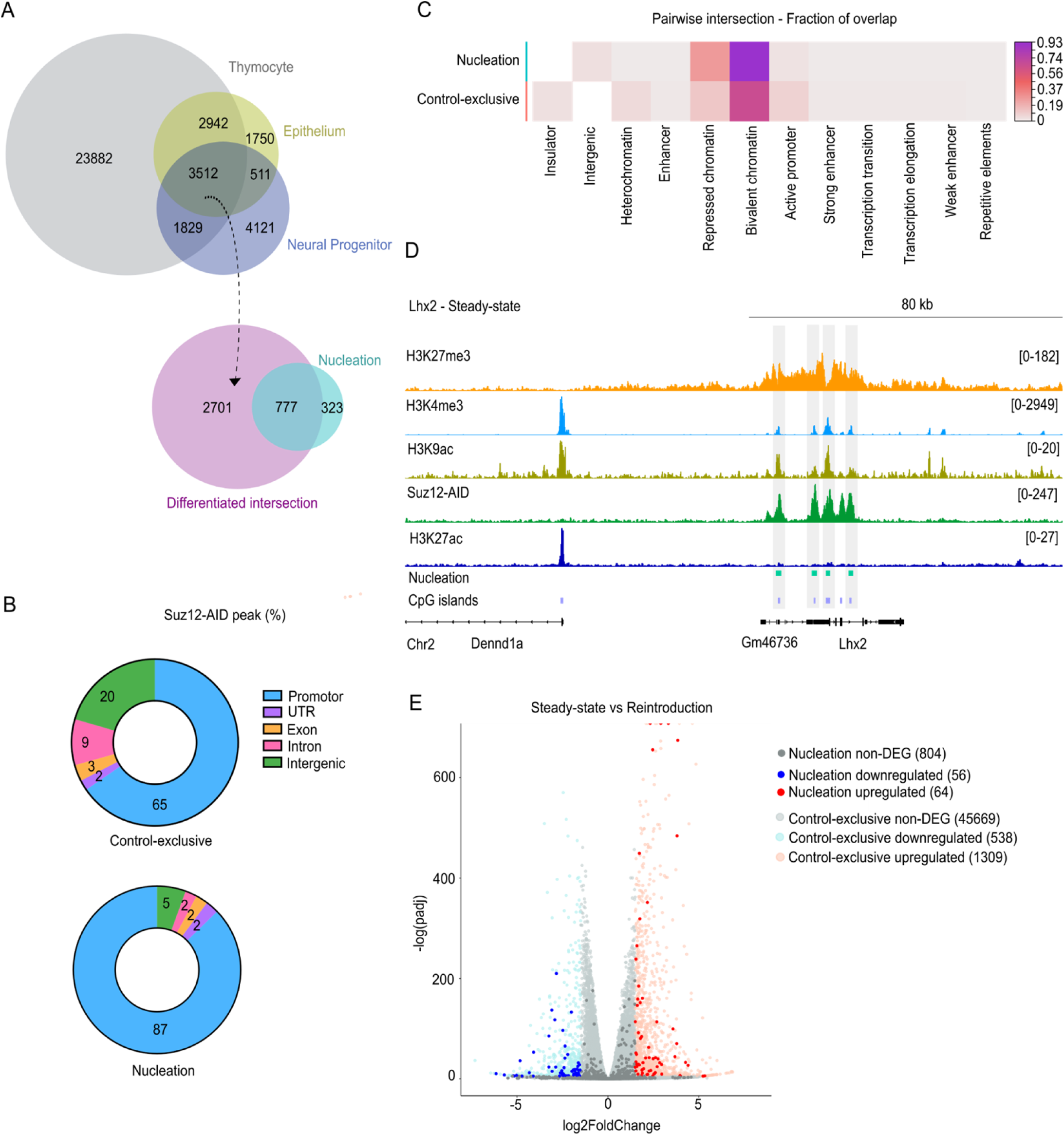
Suz12 nucleation sites are enriched at bivalent promoters. **(A)** Overlap of Suz12 ChIP-seq peaks in thymocytes (33), intestine epithelium (34), neural progenitors (35) with our mESCs nucleation sites. **(B)** Genomic distribution of Suz12 peaks across annotated genomic categories. Nucleation sites show ∼20% higher localization at promoters and reduction at intergenic regions **(C)** Chromatin state annotations from ChromHMM of mESCs. Nucleation sites are highly enriched at bivalent and repressed chromatin compared with *Control-exclusive* sites. **(D)** Representative chromatin landscape at the *Lhx2* locus at steady-state. ChIP-seq track for H3K27me3, Suz12 (this study), H3K4me3, H3K9ac and H3K27ac show bivalency at nucleation sites (gray boxes). **(E)** Volcano plot of RNA-seq differential expression (|log₂FC| > 1.5, adjusted P < 0.005) of the nearest genes to nucleation sites between *Steady-state* and *Reintroduction* conditions.

Next, we examined the genomic distribution of Suz12 nucleation sites versus *Control-exclusive* by annotating peaks for comparative analysis. In both instances, most peaks localized at promoters and intergenic regions (Fig 3B), consistent with PRC2’s known binding at gene promoters (36,37). However, nucleation sites showed ∼4-fold reduction at intergenic regions and ∼20% greater localization at promoters (Fig 3B), indicating a bias for *de novo* recruitment near transcription start sites (TSSs). To further explore the epigenetic context of these peaks, we intersected them with mESC used ChromHMM annotations (38), which classify states such as active or bivalent promoter, transcribed/elongation, strong/weak enhancer, insulator, intergenic, heterochromatin and repressed based on histone marks. Nucleation sites displayed markedly higher overlap with bivalent and repressed states (Fig 3C). Specifically, 93% of nucleation sites overlapped with bivalent chromatin (H3K4me3+H3K27me3), compared with 67% of the *Control-exclusive* peaks. Previous reports indicate that nearly 85% of H3K27me3-marked promoters are bivalent (39), but our results suggested that not all of them can function as nucleation sites. These enrichments were statistically significant when compared to a set of random peaks from the *Control-exclusive* list (Fig S3A), supporting the idea that *de novo* PRC2 recruitment is tightly associated with poised chromatin landscapes at promoters. Other studies propose that bivalency is better defined by the coexistence of H3K4me3, H3K9ac and H3K27me3 (40). To illustrate this, we visualized ChIP-seq tracks at the Lhx2 locus, where nucleation peaks coincide with the three marks and Suz12 occupancy (Fig. 3D). We also included H3K27ac, which antagonizes Polycomb deposition.

To examine the functional process associated with nucleation sites, we assessed the nearest associated gene exploring protein interactions using STRING (41). We identified 627 genes near the *de novo* recruitment peaks, forming a highly connected network enriched for pattern specification processes, and DNA-binding transcription factor activity (Fig S3B). Gene Ontology (GO) analysis of such genes revealed terms for “Cell fate commitment” (GO:0045165, FDR = 6.14E-61), “Regionalization” (GO:0003002, FDR = 7.01E-59), “Pattern specification process” (GO:0007389, FDR = 8.51E-63), and “Embryonic organ morphogenesis” (GO:0048562, FDR = 5.42E-50) (Fig S3C). Together, these data underscore the role of nucleation at bivalent promoters in safeguarding stem identity and restricting premature differentiation.

To assess the impact of Suz12 removal and reintroduction on these bivalent promoters, we conducted a transcriptional analysis. RNA-seq between *Steady-state* and *Reintroduction* revealing a total of 1,967 differentially expressed genes (DEGs; |log2FoldChange| > 1.5, P adjusted < 0.005), corresponding to ∼4% of the 48,440 annotated as genes in the mouse genome (GRCm38 M10, Ensembl 85) (Fig 3E). This is consistent with previous reports in mESCs (42). Focusing on genes nearest to nucleation sites, only 120 were DEG after *Reintroduction* (Fig 3E). Gene ontology analysis of the nucleation-associated DEGs revealed enrichment for “Forebrain development” (GO:0030900, FDR = 4.77E-10), and “Central nervous system development” (GO:0007417, FDR = 2.5E-10) (Fig S3D). These enrichments may reflect the propensity of mESCs to initiate early neurodevelopment in the absence of PRC2. Overall, these results indicate that most nucleation sites reside at bivalent loci associated with chromatin promoters and that Suz12 *Reintroduction* has relatively marginal transcriptional effects in mESCs.

### Unmethylated CGI, Polycomb cofactors and specific DNA motifs define nucleation site identity

In *Drosophila melanogaster*, Polycomb Response Elements (PREs) serve as a specific DNA sequence involved in Polycomb recruitment (43). While no direct mammalian equivalent has been clearly identified, CG-rich sequences at unmethylated CpG islands (CGI) and overrepresentation of GA/GCN tandem repeats have been associated with PRC2 targeting in mammals (14). To examine whether DNA methylation patterns correlate with nucleation sites, we analyzed the UCSC Genome Browser CGI list (44) together with a publicly available DNA methylation dataset in mESCs (45). We found that the number of unmethylated CGI is higher at nucleation sites than at *Control-exclusive* sites (Fig 4A and Fig S4A). This enrichment was statistically significant (P-value < 0.0001) when compared to the expected overlap from random sampling (Fig 4B). Moreover, nucleation sites had higher G and C nucleotide content than both *Control-exclusive* sites and random genomic regions (n = 10,000) (Fig 4C). These results are consistent with the established antagonism between Polycomb and DNA methylation and confirm that unmethylated CGI are strongly associated with *de novo* PRC2 recruitment (46).

**Fig 4.**
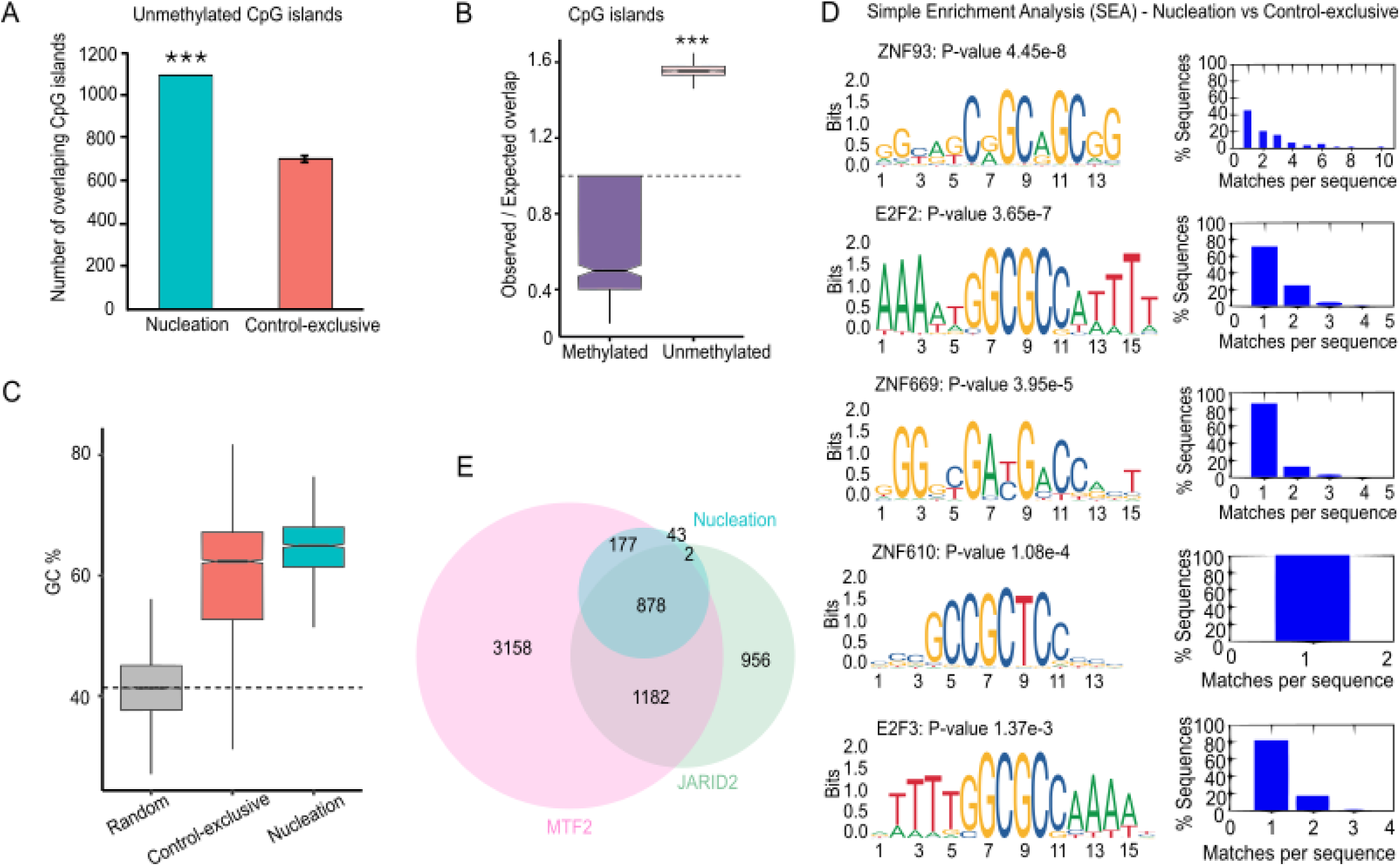
Genomic and sequence features associated with PRC2 nucleation sites. **(A)** Number of unmethylated CpG islands (CGI) overlapping with Suz12 nucleation sites or a random set of *Control-exclusive* (n = 1,000). (**B)** Observed/expected ratio of overlap between nucleation sites and methylated or unmethylated CGI, compared to random samplings of *Control-exclusive* peaks. The black dashed line marks the random expected ratio. (**C)** Cytosine and Guanine nucleotide content of random genomic regions (n = 10,000), *Control-exclusive* and nucleation sites. The dashed line marks the average GC content of the mouse genome. P-value < 0.0001 **(D)** Five selected motifs from the motif enrichment analysis of over-represented sequences that are differentially enriched at nucleation versus *Control-exclusive* sites. Matches-per-sequence profiles are shown. **(E)** Overlap between Suz12 nucleation sites with Mtf2 and Jarid2 ChIP-seq peaks in mESCs (47,48).

Then, we explored whether specific DNA motifs define our nucleation sites by applying HOMER (49) and MEME (50) motif discovery tools (S1 Table). HOMER analysis for known and novel motifs revealed matches for ZNF669, E2F6, E2F3, and E2F2 motifs. Concomitantly, MEME identified a GCC-rich motif that matches zinc-finger proteins ZNF93 and ZNF610 among the motifs differentially enriched. Further enrichment analysis using Simple Enrichment Analysis (SEA) (51) confirmed overrepresentation of the DNA binding motifs for ZNF610, ZNF669, E2F3, E2F2, and ZNF93 at nucleation sites. In Fig 4D, we present the five hit SEA motifs that were shared among studies and their Matches per sequence graphs. Overall, the identification of E2F cell cycle regulators at nucleation sites suggests a potential link to Polycomb epigenetic memory after cell division, while several zinc-finger proteins previously associated with PRC1 or PRC2 could contribute to nucleation site targeting.

Next, we examined the overlap between our nucleation sites and two well-characterized Polycomb cofactors implicated in PRC2 recruitment: the Metal Response Element Binding Transcription Factor 2 (MTF2) and Jumonji and AT-Rich Interaction Domain Containing 2 (JARID2) (52–54). By reanalyzing published ChIP-seq datasets (47,48) we found that MTF2 binding was present at 95% of our mESCs nucleation sites (1,057 sites), with a highly significant enrichment (Fisher’s test P-value < 2.2e–16) (Fig 4E). The odds ratio for MTF2 binding at nucleation versus *Control-exclusive* indicated a nearly 10-fold greater likelihood of MTF2 association at nucleation sites, supporting a strong association of MTF2 with *de novo* PRC2 recruitment. Similarly, JARID2, which has a zinc-finger domain (55), overlapped with 80% of our nucleation sites (880 sites) (Fig 4E) showing a perfect enrichment at PRC2 targets (Fisher’s test P-value = 0). Concomitantly, the odds ratio for JARID2 binding for nucleation sites versus *Control-exclusive* indicates a 16-fold higher enrichment at nucleation regions. Remarkably, 79% of nucleation sites were co-occupied by both MTF2 and JARID2, consistent with the model in which MTF2 initiates nucleation while JARID2 supports domain spreading.

Although MTF2 lacks a strict consensus motif, it preferentially binds GCG-rich, low-methylated DNA, which widens the minor groove (20,52). This relaxed helical shape of DNA is reminiscent of the DNA-RNA hybrids in R-loop structures (56), which can also stabilize PRC2-Ezh2 and PRC1-Ring1B to the chromatin (57). Thus, we sought to identify superposition between MTF2 and R-loops in mESCs by using DRIP-seq datasets (58). We found that only 11% of our nucleation sites overlap with MTF2 and R-loops (Fig S4B), with a modest enrichment (Fisher’s test, P-value > 0.05 and odds ratio of 1.75). Moreover, when comparing G-quadruplex density (a hallmark of R-loop formation), we found no statistical differences between nucleation sites and random regions (Fig S4C). These results suggest that while MTF2 recruitment to R-loops can occur, this is not a defining feature of PRC2 *de novo* recruitment. Thus, this data supports a model in which PRC2 nucleation sites correlate with unmethylated CGIs, enriched for E2F/ZNF motifs and co-occupied by MTF2/JARID2, while R-loops are not broadly required.

### Nucleation sites correlate with Polycomb-loop anchors for long-range chromatin interactions

In addition to their role in gene repression, the Polycomb proteins contribute to 3D nuclear organization by forming long-range chromatin loops (59). This prompted us to investigate whether our putative *de novo* recruitment sites are associated with such spatial interactions. We first compared the genomic distances between our Suz12 nucleation and *Control-exclusive* sites. Nucleation sites displayed a bimodal distribution: a proximal group with an interpeak distance of <10 kb interpeak distance, and a distal group spanning nearly 8 Mb (Fig 5A and S5A). The proximal group includes 122 cases that span multiple nucleation sites within a gene (e.g., Barx1, Nkx2-2, and Foxa2). By contrast, the median interpeak distance for *Control-exclusive* sites was ∼100 kb, whereas nucleation peaks were spaced ∼1 Mb apart, with some exceeding 10 Mb (Fig 5A). The 10-fold increase in spacing at nucleation sites suggests that they may be involved in longer-range chromatin interactions. To explore this possibility, we analyzed histone H3K27me3 HiChIP-seq data from mESCs (59) at 10 kb resolution to capture both short and long interactions. We identified 4,254 H3K27me3-mediated interactions, of which 260 were mediated by one nucleation site and an average distance of 0.33 Mb, and 102 were anchored by two nucleation sites with an average distance of 0.86 Mb (Fig 5B). Thus, the average nucleation anchor-mediated interaction does not exceed 1 Mb. Among these, the longest loop spanned 7.58 Mb, connecting nucleation sites at Barx1 and Neurog1. All remaining interactions with non-nucleation anchors have an average interaction distance of 0.18 Mb. According to a one-way ANOVA test, the interactions mediated by two-nucleation anchor interactions are significantly longer (P value of < 0.0001) than those intervened by one-nucleation or non-nucleation anchors (Fig 5C).

**Fig 5.**
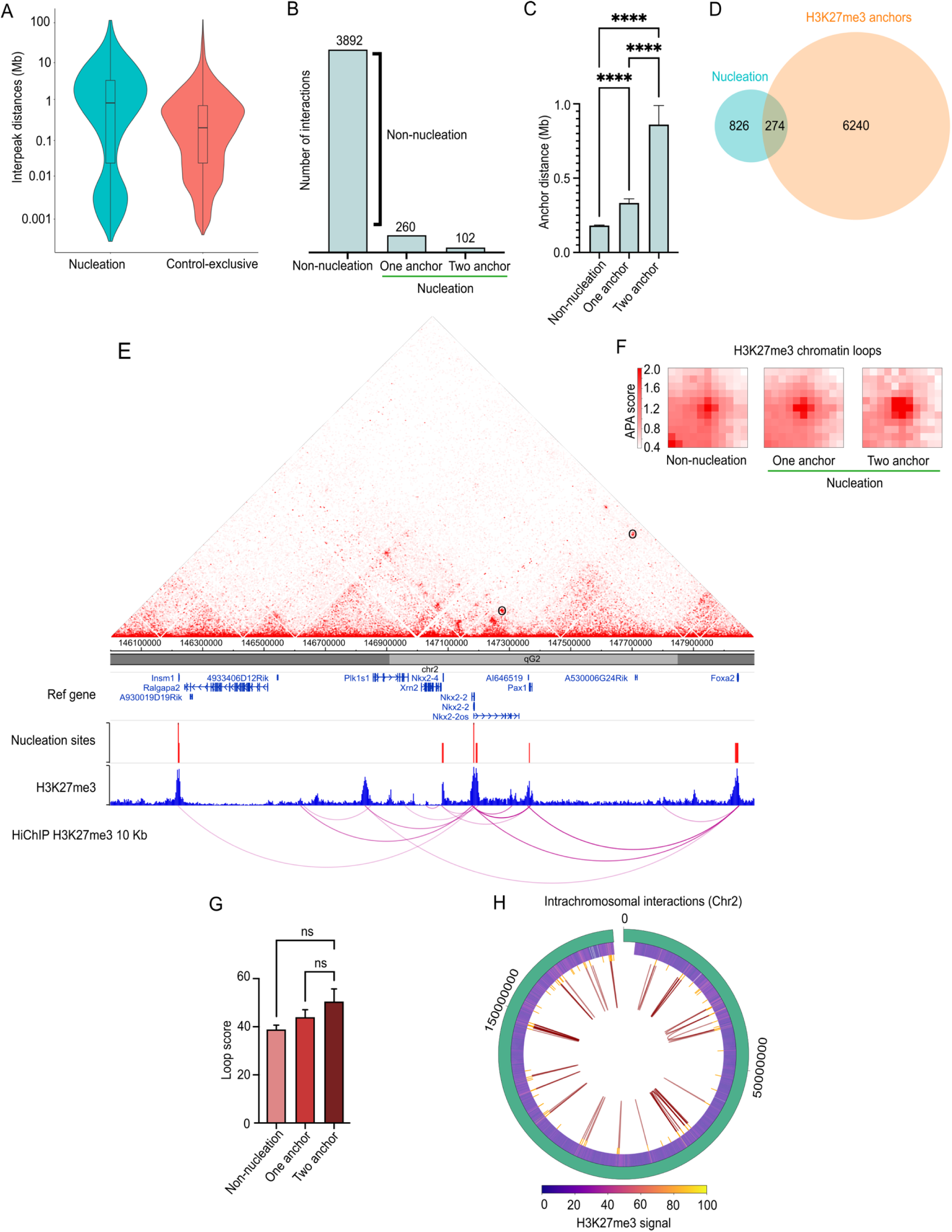
Nucleation sites act as anchor points for H3K27me3-mediated chromatin loops. (**A**) Interpeak distances of nucleation and *Control-exclusive* Suz12 peaks. Nucleation sites exhibit significantly longer distances. (**B**) Global interaction counts involving one or two nucleation sites as loop anchors based on histone H3K27me3 HiChIP data (59). (**C**) Anchor nucleation distances. One-way ANOVA test of P-value of < 0.0001. (**D**) Overlap between nucleation sites and H3K27me3 loop anchors. (**E**) HiChIP contact matrix (10 kb resolution) showing H3K27me3-associated interactions for the Insm1-Nkx2-Foxa2 region in mESCs. ChIP-seq tracks for histone H3K27me3 (blue), nucleation site (red) and its H3K27me3-mediated loops seen as a virtual 4C assay. Loops identified on the Hi-ChIP matrix are shown as black circles. (**F**) Aggregate Peak Analysis (APA) of the H3K27me3 Hi-ChIP comparing the interaction strength for loops without nucleation anchors, with one anchor, or two nucleation anchors at the Chr2 (n = 102 sites). **(G)** One-way ANOVA test of loop scores. P value of < 0.05. (**H**) Circos plot of Chr2 with intrachromosomal H3K27me3 interactions (red loops) mediated by at least one nucleation site (yellow) as anchor point. Interactions occur primarily in the parts of the genome that have histone H3K27me3 enrichment (middle ring).

Furthermore, we found that approximately 25% of our Suz12 nucleation sites overlap with these histone H3K27me3 HiChIP-seq peaks (Fig 5D), suggesting a coupling of *de novo* PRC2 recruitment to high-order genome architecture. To investigate the spatial features of these loops in more detail, we built contact maps around anchor nucleation sites, integrating HiChIP contact matrix, nucleation coordinates, histone H3K27me3 enrichment peaks, and virtual 4C representation of chromatin interactions. These contact maps revealed overlap between histone H3K27me3 peaks, nucleation sites, and loop anchor points. For example, consistent with previous report (14), nucleation sites near Evx2 formed long-range interactions within the vicinity of the HoxD locus and other distant nucleation sites (Fig S5B). When centered on Pax1 as viewpoint, significant interactions were observed leading to Foxa2 (0.67 Mb) and Nkx2-2 (0.17 Mb) loci (Fig 5E), two of the strongest interactions, as indicated by the thickness of the loop arcs. We further noticed that H3K27me3 signal intensity at *Steady-state* is higher at nucleation-anchored loops compared to non-nucleation mediated loops (Fig S5C). Therefore, we infer that nucleation-anchored loops act as hubs that concentrate PRC2/H3K27me3, facilitating mark spreading between linked loci.

To quantify interaction strength, we performed Aggregate Peak Analysis (APA), which quantifies the strength and significance of chromatin loops. When comparing the same number of H3K27me3 loops genome-wide (n = 102), we noticed a higher APA score at those with nucleation sites as anchors compared to the loops formed without nucleation sites (Fig 5F). In particular, the loops with two nucleation anchors are the most robust, exhibiting a concentrated signal with defined contrast and a focalized pattern. However, a one-way ANOVA test did not reach statistical significance (Fig 5G), although nucleation-anchored loops showed a consistent trend toward higher score. Hence, the average intensity of chromosomal interactions is higher when two nucleation anchors mediate the loops. The robustness of these loops suggests a stable chromosomal organization that likely plays a crucial role in gene regulation and the maintenance of repressive chromatin domains. The chromosomes (Chr) with the most nucleation sites acting as looping anchors are Chr1 and Chr2. This finding is consistent with previous evidence highlighting the organizational role of Polycomb in nuclear architecture, particularly on chromosomes with large regulatory gene clusters, such as Chr2 and Chr17 (60,61). To visualize this, we generated a circos plot of Chr2 intrachromosomal interactions mediated by at least one nucleation site (Fig 5H). We identified 54 H3K27me3-loops at Chr2 which supports the concept of sub-megabase Polycomb interacting neighborhoods. In sum, nucleation sites preferentially anchor and strengthen Polycomb loops, linking *de novo* PRC2 recruitment to high-order genome architecture

## Discussion

In this work we identified early nucleation sites of PRC2 in mouse embryonic stem cells (mESCs) using an inducible and reversible degradation system for Suz12, a core component essential for assembling the complex and its recruitment (1). This approach enabled us to track early stages of *de novo* Polycomb recruitment genome-wide, revealing over 1000 high-confidence putative nucleation sites. Our findings suggest that *de novo* PRC2 recruitment is linked to a combination of chromatin states, DNA features, cofactor recruitment, and spatial genome organization (Fig. 6).

**Fig. 6.**
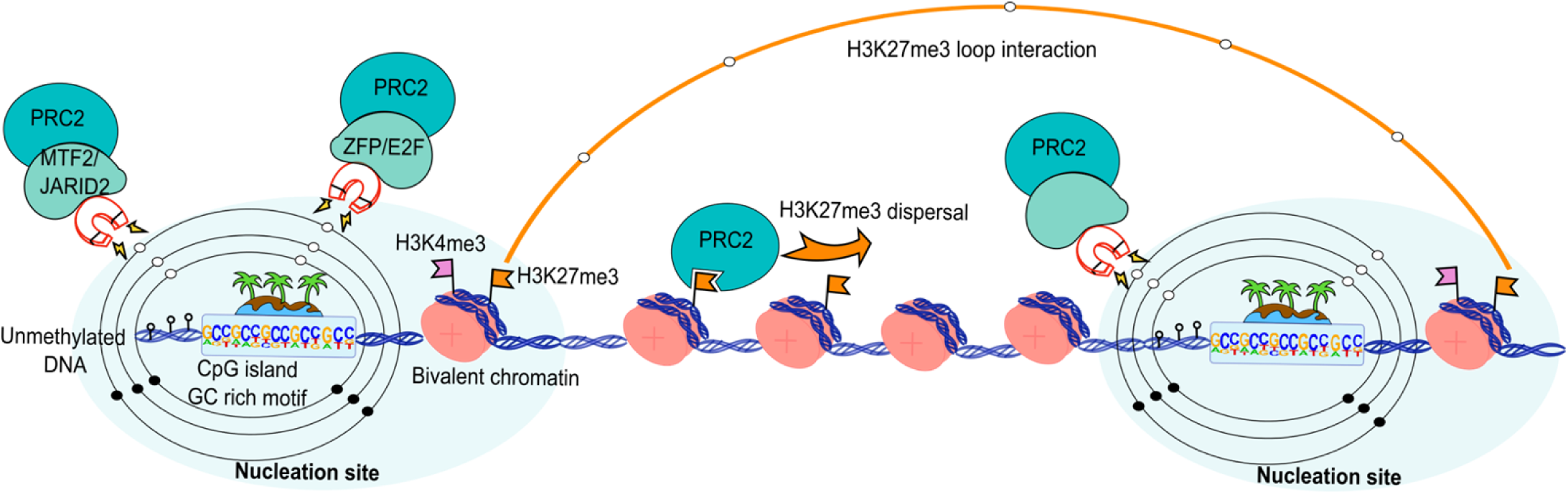
Graphical abstract of features associated with PRC2 recruitment. Nucleation sites act as magnets for *de novo* PRC2 recruitment. Potential recruiting partners such as MTF2, JARID2, zinc-finger proteins (ZFPs), and E2F cell cycle regulators are hallmarks of these nucleation sites. Their attractiveness correlates with the presence of unmethylated CGI at gene promoters, characterized by bivalent chromatin. The histone H3K27me3 disperses from nucleation sites while they can interact with one another, strengthening long-distance loop interactions.

We found that after depletion and reintroduction of Suz12, only modest transcriptional changes occurred, supporting the view that *de novo* recruitment in mESCs primarily establishes a poised chromatin state rather than actively repressing transcription. This functional poising likely preserves developmental potential, allowing rapid gene activation (or silencing) upon differentiation, and is consistent with previous findings that PRC2 loss does not compromise self-renewal but may impair differentiation (62,63). Remarkably, nearly 90% of the nucleation sites are located at bivalent promoters and are approximately twice as broad compared to *Control-exclusive* peaks, likely reflecting condensation of Polycomb complexes during early recruitment (64,65). Developmental regulatory genes are frequently marked by this bivalent state, where H3K27me3 and H3K4me3 coexist. Such genes are typically expressed at low levels in mESCs to undergo rapid activation or repression upon differentiation. Interestingly, ∼70% of these sites are shared across tissues from all three germ layers, suggesting a universal role in developmental gene regulation, whereas the remaining (30%) sites appear stem-cell-specific and may resolve during lineage commitment.

Shortly after Suz12 reintroduction, nucleation sites show strong Suz12 and histone H3K27me3 gains, supporting the notion that these sites act as binding hubs. We found that nucleation events highly correlate with unmethylated CGI and exhibit higher local CG content, consistent with the antagonism between DNA methylation and Polycomb occupancy. This epigenetic context appears to enhance PRC2 accessibility and stabilization, in part via interaction with accessory partners such as MTF2 and JARID2, as the ChIP-seq dataset overlapped 95% and 80%, respectively, with our nucleation sites. JARID2 facilitates PRC2 recruitment by PRC1-mediated H2AK119ub1 recognition, supporting localized PRC2.2 *de novo* recruitment that relies on the CGG and GA sequence motifs (54). Although JARID2 can restore Suz12 binding in the absence of MTF2 (14), it does so with slower kinetics. Conversely, most evidence strengthens the idea that MTF2 ensures PRC2 recruitment at Polycomb key genes, whereas JARID2 is essential for dispersing new H3K27me3 domains (13). Thus, these chromatin-binding proteins play essential roles in domain nucleation, significantly enhancing their binding chances to *de novo* recruitment sites compared to other PRC2 targets. Nevertheless, additional interactions with other protein partners or RNA molecules may contribute to nucleation, such as lncRNAs or chromatin-associated RNAs that have been demonstrated to play an important role in Polycomb targeting (66–70).

At sequence level, nucleation sites are enriched for GCC/GC-rich motifs. While Polycomb Response Elements (PREs) can specify targeting in invertebrates (43,71), in mammals, the only genomic signature associated with Polycomb recruitment is the enrichment of GC-rich sequences at unmethylated CGI promoters, often accompanied by GA or GCN tandem repeats (14,72,73). Within this CG-rich context, our motif analysis was resolved two major classes: E2F cell cycle regulators and zinc-finger proteins (e.g. ZNF93, ZNF610 and ZNF669), suggesting a sequence-encoding docking for PRC2 cofactors. In particular, E2F2/3 are activators of G1-S genes and are critical for cell cycle re-entry following arrest (74,75). Their enrichment at nucleation sites may reflect a mechanism to orchestrate Polycomb-epigenetic memory after cell division. While evidence for E2F-mediated recruitment of PRC2 is limited, E2F6 can act as an accessory protein of PRC1 complexes in quiescence and germline gene silencing (75–77), and it has also been associated with PRC2 during silencing of meiotic genes (76). However, only a small fraction of Suz12 co-localizes with E2F6 (77). Regarding the zinc-finger proteins, ZFP277 interacts with PRC1 through the PCGF4 subunit at the Ink4a/Arf locus (78). Also, POGZ associates with PRC1 during neuronal differentiation (79), while ZFP217 and ZFP516 indirectly interact with PRC2 through the transcriptional corepressor Ctbp2 during mESCs differentiation (80). These examples are consistent with the notion that zinc-finger proteins could participate in PRC2 recruitment (81). Together, these observations support a model in which clustered GC/GCC motifs and specific TFs (E2Fs, ZNFs) recruit Polycomb cofactors to a subset of CGI promoters, facilitating PRC2 nucleation and H3K27me3 spreading; future experiments editing these motifs at endogenous loci, or reconstituting them ectopically, will define their involvement in PRC2 nucleation and their roles in maintaining the epigenetic landscape.

Beyond the DNA sequences and chromatin context, we observed that PRC2 nucleation sites correlate with a spatial network of long-range chromatin loops ranging from 1 Mb to 10 Mb, which are significantly longer than the average H3K27me3 loop distance in *Control-exclusive* conditions (∼100 kb). These long-range interactions among nucleation sites are more robust when both anchor points are nucleation sites. However, only about 25% of nucleation sites overlapped with H3K27me3 HiChIP-detected loop anchors. This observation suggests that only a subset of nucleation sites form long-range contacts that may seed spreading of H3K27me3 in *cis* and in *trans*.

Finally, our findings must be interpreted considering models that emphasize the role of PRC1 in initiating Polycomb recruitment and long-range interactions (8). In particular, RING1B-mediated monoubiquitylation of H2AK119 has been shown to promote PRC2 binding (54,82). Future studies should also explore how PRC1 architecture and activity intersect with PRC2 nucleation and memory.

## Concluding remarks

Our data indicate that early PRC2 nucleation in mESCs are concentrated at broad, GC-rich bivalent promoters, frequently co-occupied by MTF2 and JARID2, and embedded within a subset of H3K27me3-linked long-range interactions. These features -chromatin state, sequence composition, cofactor occupancy, and 3D architecture-provide a coherent framework for how PRC2 re-establishes its occupancy after domain erasure, with limited transcriptional impact at early time points and with many sites conserved across germ-layer lineages. The predominance of nucleation at bivalent promoters highlights how this chromatin state serves as a privileged entry point for Polycomb targeting, balancing transcriptional flexibility with repressive stability. While these associations are compelling, they are correlative. Definitive tests will require motif disruption and transplant experiments at endogenous loci, timed depletion/restore of MTF2 and JARDI2, and dissection of PRC1 contributions. Mapping the order and dependency of these events should clarify how nucleation sites act as hubs to initiate local deposition and promote spreading within repressive neighborhoods. By focusing on the onset of d*e novo* recruitment rather than steady-state profiles, this work outlines sequence and chromatin encoded entry points that can be functionally interrogated to explain how Polycomb domains, and the landscape they embody, operate to acquire and maintain a stable epigenetic memory.

## Methods

### Cell culture and auxin treatment

J1 mouse embryonic stem cells (mESCs) were cultured under standard conditions on 0.1% gelatin-coated plates and in ES medium: DMEM High Glucose supplemented with 15% FBS (Biowest L0101-500), 0.1 mM 2-mercaptoethanol, 2 mM L-glutamine, 0.1 mM MEM non-essential amino acids (NEAA), 1% nucleoside mix, 50 U/mL Penicillin/Streptomycin (P/S), Leukemia inhibitory factor (LIF, made in house) and 2i inhibitors: 1 μM PD0325901 and 3 μM CHIR99021. Cells were passed every other day using 0.05% Trypsin/EDTA. HEK293T cells were used for lentiviral production in DMEM supplemented with 10% FBS, 2 mM L-glutamine, 0.5 mM sodium pyruvate, and 50 U/mL P/S. All cells were grown at 37°C in a humidified incubator with 5% CO₂. Auxin (indole-3-acetic acid, IAA; Sigma, catalog no. I5148) was dissolved in water to a 200 mM stock solution and stored at -20°C. For AID-mediated degradation, auxin was titrated and used at a final concentration of 50 μM in culture media.

### Cloning

To generate the CRISPR donor plasmid, four fragments (5’ coding and 3’ UTR homology arms, mAID-mClover-NeoR cassette, and pBluescript backbone) were amplified with Phusion High-Fidelity DNA polymerase (ThermoScientific) using Gibson assembly with the In-Fusion kit (Takara). Homology arms were amplified from mESCs genomic DNA. The mAID-mClover-NeoR sequence was subcloned from pMK289, from Masato T. Kanemaki’s laboratory (83), and the backbone from pBluescript SK (-). To prevent recleavage by Cas9, we disrupted the PAM sequence within the donor template. The gRNA was selected using Benchling (https://benchling.com/faq) and cloned by Gisbon assembly into pSpCas9(BB)-2A-Puro (PX459) V2.0 (Addgene 62988;). The pRAIDRS-P7 NLS-mOrange-AID from Ran Brosh (26) was modified by Gibson to exclude the AID sequence. Primer sequences are in the S1 Table. All plasmids’ constructs were propagated in *Stbl3* competent cells One Shot Stlb3^TM^ (Invitrogene) and NEB®Stable Competent *E. coli* (New England).

### Suz12 Auxin-Inducible Degradation knock in

mESCs were co-transfected with the gRNA plasmid and the donor plasmid (2:1) using Polyethylenimine (PEI 1:3) and selected with neomycin (1.8 ug/mL). mClover cells were sorted as GFP+ using a BD FACSMelody Cell Sorter. Single mESCs colonies with typical morphology were manually picked for genotyping by PCRs using primers surrounding the target site along with Sanger sequencing verification of in-frame insertion. Clones with correct insertion were validated by western blot and transduced with lentiviral particles to integrate the OsTIR1 receptor; lentiviral particles were generated co-transfecting the pRAIDRS-P7 NLS-mOrange-notAID along with psPAX2 and pMD2.G using PEI; particles were concentrated using Amicon Ultra-15 (Millipore). Double positive GFP/Orange cells were sorted and the integration of OsTIR1 was verified by PCR genotyping. After auxin treatment, cells were fixed with ice-cold paraformaldehyde at 4% and the GFP median fluorescence was measured by FACS.

### Western blot

Cells were lazed in RIPA buffer (1% NP-40, 0.5% deoxycholate, 0.1% SDS, 150 mM NaCl, 50 mM Tris-HCl pH 8, with Protease Inhibitor Cocktail, Roche) and incubated on ice for 30 minutes with occasional vortexing. Lysates were centrifugated at 14,000 x g for 15 minutes at 4°C and protein content was determined on the supernatant. For histone extraction, 1×10^7^ cells we resuspend in a Triton Extraction Buffer (PBS, 0.5% Triton X-100, 2 mM PMSF, 0.02% sodium azide) followed by a 10-minute centrifugation at 400 x g at 4°C, pellet was resuspended in 0.2 N HCl (4 x10^7^ cells/mL). Histones were acid extracted overnight at 4°C. Protein content was measured with DC Protein Assay Reagents (BioRad, 5000116). 20–40 µg of total histones was resolved by SDS–PAGE and transferred to PVDF (0.45 μm, Immobilon-P) or nitrocellulose (0.2 um, Amersham) membranes. Membranes were blocked with TBST (10 mM Tris-HCl pH 7.9, 150 mM NaCl and 0.05% Tween-20) with 5% skim milk, incubated overnight with primary antibodies and washed three times with TBST. HRP-conjugated secondary antibodies were applied, and blots were developed with Immobilon HRP substrate and imaged using a C-DiGit Blot Scanner (LI-COR) with minor adjustments. Antibodies: rabbit anti-GFP (Abcam, ab290, 1:2000), rabbit anti-SUZ12 (Thermo Fisher, 39357, 1:1500), rabbit anti-H3K27me3 (Millipore, 07449, 1:1000), rabbit anti-H3 (Abcam, ab1791, 1:10,000), mouse anti-Actin (Abcam, ab197277, 1:10,000), mouse anti-GAPDH (Sigma, G8795, 1:5000), goat anti-rabbit (Santa Cruz, sc2030, 1:10,000) and goat anti-mouse (Santa Cruz, sc2031, 1:10,000).

### Chromatin immunoprecipitation (ChIP)

Protocol was performed as described Lee et al. (84,85) with modifications: cells were dissociated with 0.05% trypsin, fixed with 1% formaldehyde solution for 10 min at RT, and quenched with fresh 125 mM glycine for 5 min RT. After PBS washed, nuclei were isolated using sequential lysis buffers: B1 (50 mM HEPES-KOH, pH 7.5, 140 mM NaCl, 1 mM EDTA, 10% Glycerol, 0.5% NP-40, 0.25% Triton X-100, 1x protease inhibitors; 10 min at 4°C), B2 (10 mM Tris-HCL, pH 8, 200 mM NaCl, 1 mM EDTA, 0.5 mM EGTA, and 1x protease inhibitors; 10 min at 4°C), and B3 (10 mM Tris-HCL, pH 8, at 4°C, 100 mM NaCl,1 mM EDTA, 0.5 mM EGTA, 0.1% Na-deoxycholate, 0.5% N-Lauroylsarcosine sodium salt, and 1x protease inhibitors). Chromatin was fragmented to an average size of 250-500 bp using a Diagenode Bioruptor and Triton X-100 was added to a final concentration of 1%. ChIP was performed using 1 ug of antibody per 2 million cells incubated and rotated overnight at 4°C. Antibodies: rabbit anti-GFP (Abcam, ab290), rabbit anti-H3K27me3 (Cell signaling, 9733). Dynabeads (10 µL pero µg Ab) were pre-washed in PBS + 0.5% BSA and rotated for 2 hours at 4°C before adding to the IPs. Then washes X5 with ChIP RIPA buffer 4°C (50mM HEPES-KOH, pH 7.5 at 4°C, 500 mM LiCl, 1 mM EDTA, 1% NP-40, and 0.7% Na-deoxycholate) and one wash with TE + 50 mM NaCl. DNA was eluted in freshly prepared elution buffer (50 mM Tris, pH 8, 10 mM EDTA, 1% SDS) at 65°C for 20 min and then de-crosslinked overnight at 65 °C. Samples were treated with RNase A and Proteinase K and DNA was purified by phenol:chloroform:isoamyl alcohol (P:C:IA). Libraries were prepared using NEBNext Ultra II DNA Library Prep Kit (NEB), quantified by Qubit dsDNA HS Assay, quality-checked with Aligent Bioanalyzer HS DNA chip. Libraries were sequenced as 150 pb paired-end reads on the Illumina HiSeq platform.

### RNA extraction and sequencing

Total RNA was purified in triplicates from mESCs with TRIzol (Life Technologies), and RNA integrity (RIN) was corroborated using an Aligent Bioanalyzer. We follow the manufacturer’s TruSeq RNA Sample Prep v2 (Illumina, USA) instructions for transcriptome sequencing. The mRNA was purified utilizing poly-T oligo-attached magnetic beads. After purification, the mRNA was fragmented and converted into first-strand cDNA utilizing reverse transcriptase and random primers. The second strand, cDNA, is then synthesized using DNA Polymerase. Actinomycin D was added to prevent DNA-dependent synthesis and improve strand specificity. The cDNA fragments undergo a single ’A’ base addition and subsequent adapter ligation. After purification and PCR enrichment, the final cDNA library is suitable for subsequent cluster generation and DNA sequencing. We sequenced the libraries using a NextSeq 500 platform (Illumina) in 75 bp paired end reads format.

### Preprocesing and maping of Next-generation sequencing data

All Raq FASTQ files generated in this study or retrieved from public repositories were processed using a uniform pipeline: Adapter sequences and low-quiality reads were trimmed with Trim Galore v0.6.10 with default parameters (86). Files were verified with FastQC v0.12.1 (87). Trimmed reads were aligned to the mm10 mouse reference genome using Bowtie2 v2.5.3 then BAM files were obtained using samtools 1.20; *samtools view -bS*, then sorted and indexed with *samtools sort* and *samtools index* with default parameters (88,89).

### ChIP-Seq data analysis

Our ChIP-Seq data and public dataset BAM files were processed using a standardized pipeline. Peak calling was performed per replicate using MACS2 v2.2.7.1; *callpeak* with default parameters (90). The peaks obtained from the processing of our auxin induced experiment were merged into non-redundant union sets for each of the immunoprecipitation antibodies with Bedtools v2.30.0; *bedtools merge* with default parameters (91). We then used the unified peak sets to count reads that overlapped with each peak using FeatureCounts v2.0.6 (92). We then used the counts to build a matrix suitable for its use with DESeq2 v1.38.0 within R v4.2.3 environment, example scripts with our utilized conditions and parameters publicly available (https://github.com/cperalta22/suz12_nucleation_mesc) (93). Plots for downstream visualization were generated with ggplot2 3v.4.2. To compare and intersect BED files obtained from the peak calling of our samples and re-analyzed public datasets as well ChromHMM annotation were assessed using the command *intervene venn* with the parameter *–save-overlaps* from the package Intervene v0.6.5 (94). Fisher’s exact tests were performed with BEDtools v2.30.0 using *bedtools fisher* with default parameters. Peak annotation and visualization were conducted using ChIPseeker (Galaxy v1.28.3+galaxy0) (95). To create visualization files and plots such as BigWig and heatmaps, we calculate scaling factors for all samples for each immunoprecipitation condition with deepTools 3.5.0; *multiBamSummary* with *–scalingFactors* as a parameter, we used the normalization factors as a parameter of deepTools using *bamCoverage –*scaleFactor *-v –extendReads –binSize 5*. We calculated and created new BigWig files for every plot that required the usage of a different set of BigWig files as input. To obtain signal heatmaps we utilized deepTools using *computeMatrix and plotHeatmap.* Comparative bigwig tracks were created with bigwigCompare from the deeptools suite, taken as input the normalized and scaled merged bigwig files for each experimental condition evaluated. Gene ontology and protein interaction analysis were performed with STRING v12.0 (41). Motif analysis was assessed with MEME suite (v5.5.7) identification tools (96), using Motif Analysis of Large Nucleotide Datasets (MEME-ChIP) (50) and Simple Enrichment Analysis (SEA 5.5.7) (51), both with default parameters. HOMER Motif Discovery and Analysis (49) was assessed with suite (v5.1) with default parameters.

### Differential gene expression analysis

With sorted BAM files as input we estimated raw sequencing reads abundance over the Ensembl gene annotation (mm10) mouse genome using FeatureCounts v2.0.6. Then with DESeq2 1.38.0 running under an R 4.2.3 environment we estimated gene expression differences. Scripts with our utilized conditions and parameters are available at: https://github.com/cperalta22/suz12_nucleation_mesc. Plots for downstream analysis were made using ggplot2 v3.4.2

### Code and data availability

Raw sequencing data generated in this study is available via the Gene Expression Omnibus accession number: ChIP-seq GSE305054 and RNA-seq GSE305055, in addition we utilized publicly available datasets under the following accession numbers: thymocytes GSM1498452 (33), intestinal epithelium GSM3020554 (34), neural progenitors GSM878558 (35), H3K4me3 GSM6261533 (97), H3K9ac GSM8107956 (98), H3K27ac GSE280487 (99), Mtf2 GSM6585902 (47), Jarid2 GSM6585904 (48), R-loops GSM1720620 (58), HiChIP-seq GSE150907 (59), and DNA methylation GSE266926 (45). Scripts with the code and full parameter list of the commands mentioned on this methods section are available on the following repository: https://github.com/cperalta22/suz12_nucleation_mesc. In addition, we forked the original repository (https://github.com/guifengwei/ChromHMM_mESC_mm10) containing the BED files for the ChromHMM annotation of mESCs, fork is available on the following link: https://github.com/cperalta22/ChromHMM_mESC_mm10.

### Random distribution analysis

A set of random peaks was selected from a list of background (control exclusive) peaks using a perl script (v. 5.34) with the random function. The number of random peaks selected was equal to the number of nucleation sites. The number of random peaks that overlap with different chromatin states was assessed with intersectBed from bed tools (v2.31.1) using the -wa option. The enrichment of G-quadruplexes within random peaks and nucleation sites was calculated using computeMatrix in scale-regions mode from deeptools (v3.5.6). The process was repeated 1,000 times with a bash script to get a random distribution. The resulting random distributions were compared to the distribution of nucleation sites and plotted in R (v 4.5.0) with ggplot2 (v 3.5.2). The coordinates for chromatin states were obtained from https://github.com/guifengwei/ChromHMM_mESC_mm10 and the enrichment for G-quadruplexes from GSM5259790 (100).

### CpG methylation and CG percentage

A table with single-base pair methylation levels derived from control mESCs was downloaded from GSE266926 (45). Only sites with >50% methylation were considered as methylated. A bed file of CpG islands (CGI) was downloaded from the UCSC genome browser table (mm10 version). The number of methylated CpGs at CGI was calculated by overlapping the CpG bed file with the mESCs methylation file intersectBed using the -c option. Islands with > = 20% of methylated CpGs were considered as methylated, and islans with <20% of methylated CpGs were considered unmethylated. Methylated and unmethylated CGI were overlapped with nucleation sites or a random set of peaks selected from control exclusive (1,100 peaks). The randomization was repeated 1,000 times. Observed/Expected values were obtained by dividing the values of nucleation / random peaks. For CG percentage, the fasta sequence from nucleation, control exclusive, and a set of 10,000 random peaks (obtained with bedtools random) of the same size as nucleation sites was obtained. GC percentage was calculated with an in-house perl script. All results were plotted in R (v 4.5.0) with ggplot2 (v 3.5.2).

### HiChIP analysis

Histone H3K27me3 HiChIP data from mouse embryonic stem cells (mESCs) (59). The .hic file was converted into a multi-resolution .mcool file using hic2cool utility (https://github.com/4dn-dcic/hic2cool). Chromatin loop detection was performed at 10 kb resolution using pyHICCUPS script from hicpeaks (101) package with default parameters. The resulting BEDPE file was intersected with a BED file containing nucleation site coordinates. Aggregate Peak Analysis (APA) was carried out using the apa-analysis script from hicpeaks (101). Visualization of HiChIP contact matrices at selected genomic regions was performed using the tadlib package (102,103). A circos plot showing H3K27me3 loops on chromosome 2 was carried out using pyCirclize (moshi4. (n.d.). and vizualized in Python (Retrieved July 1, 2025, from https://moshi4.github.io/pyCirclize/) (104).

## Acknowledgments

Itzel Alejandra Hernández-Romero conducted this study to fulfill the requirements of Programa de Doctorado en Ciencias Bioquíımicas of Universidad Nacional Autónoma de Mexico (UNAM), and received a doctoral scholarship from Secretaría de Ciencia, Humanidades, Tecnología e Innovación (SECIHTI, #CVU 886138). We thank the technical assistance of Beatriz Aguirre López for in-house rLIF production and José Fernando Becerra-Vélez for technical advice. At IFC, we thank the Taller, Biblioteca, Bioterio and Unidades de Biología Molecular e Imagenologia: Laura Ongay-Larios, Guadalupe Codiz, Minerva Mora and Ruth Rincón. We also thank Ran Brosh for pRAIDRS system plasmid donation (26), and Masato T. Kanemaki for the pMK289 (mAID-mClover-NeoR) plasmid donation (83).

## Author Contributions

Conceptualization: IAHR, and VJV; Data curation: IAHR and CAPA; Formal analysis: IAHR, CAPA, ARF and DVG; Funding acquisition: VJV, JW, and FRT; Investigation: IAHR and VJV; Methodology IAHR and MSN; Project Administration: VJV, JW, NCD and HZ; Resources: HZ, NCD, MSN, MFM, and SAR; Software: IAHR, CAPA, ARF, ACPH and DVG ; Supervision: VJV, JW, and FRT; Visualization: IAHR, CAPA, DVG and ARF; Writing – original draft: IAHR, and VJV; Writing – review & editing: IAHR, VJV, JW, CAPA, ARF, FRT, SAR, MFM. All authors have read and approved the published version of the manuscript.

## Copyright

2025 Hernández-Romero et al. This open-access article is distributed under the terms of the Creative Commons Attribution License, which permits unrestricted use, distribution, and reproduction in any medium, provided the original author and source are credited.

